# The TRIM9/TRIM67 neuronal interactome reveals novel activators of morphogenesis

**DOI:** 10.1101/2020.10.02.323980

**Authors:** Shalini Menon, Dennis Goldfarb, Tsungyo Ho, Erica W. Cloer, Nicholas P. Boyer, Christopher Hardie, Andrew J. Bock, Emma C. Johnson, Joel Anil, M. Ben Major, Stephanie L. Gupton

## Abstract

TRIM9 and TRIM67 are neuronally-enriched E3 ubiquitin ligases essential for appropriate morphogenesis of cortical and hippocampal neurons and fidelitous responses to the axon guidance cue netrin-1. Deletion of murine *Trim9* or *Trim67* results in neuroanatomical defects and striking behavioral deficits, particularly in spatial learning and memory. TRIM9 and TRIM67 interact with cytoskeletal and exocytic proteins, but the full interactome is not known. Here we performed the unbiased proximity-dependent biotin identification (BioID) approach to define TRIM9 and TRIM67 protein-protein proximity network in developing cortical neurons and identified neuronal putative TRIM interaction partners. Candidates included cytoskeletal regulators, cytosolic protein transporters, exocytosis and endocytosis regulators, and proteins necessary for synaptic regulation. A subset of high priority candidates was validated, including Myo16, Coro1A, SNAP47, ExoC1, GRIP1, PRG-1, and KIF1A. For a subset of validated candidates, we utilized TIRF microscopy to demonstrate dynamic colocalization with TRIM proteins at the axonal periphery, including at the tips of filopodia. Further analysis demonstrated the RNAi-based knockdown of the unconventional myosin Myo16 in cortical neurons altered axonal branching patterns in a TRIM9 and netrin-1 dependent manner. Future analysis of other validated candidates will likely identify novel proteins and mechanisms by which TRIM9 and TRIM67 regulate neuronal form and function.

## INTRODUCTION

Neuronal morphogenesis and function require coordinated cytoskeletal reorganization and plasma membrane expansion (McCormick and Gupton, 2020). Exocytosis leads to the addition of plasma membrane (Pfenninger, 2009; Gupton and Gertler, 2010; Winkle *et al*., 2014; Urbina and Gupton, 2020), whereas cytoskeletal reorganization molds the plasma membrane into the complex arborized neuronal shape. The functions of many cytoskeletal and membrane remodeling proteins are defined, yet how their functions are coordinated to promote cell shape change remains poorly understood. Further there is likely specialized machinery for the unique cell shape changes that occur in neurons, which achieve an elongated and polarized morphology in response to a host of guidance cues. Phylogenetic analysis suggested that the emergence of the class I TRIM (TRIpartitite Motif) E3 ubiquitin ligases and the emergence of neuronal-like cells appeared to have occurred contemporaneously (Boyer *et al*., 2018), and thus class I TRIMs may be well poised to regulate neuronal form and function.

The TRIM family is a subclass of RING-domain E3 ubiquitin ligases, which comprises over 80 human members, some of which are brain-enriched and implicated in neuronal development and neurological disorders (Tocchini and Ciosk, 2015; Jin *et al*., 2017; Watanabe and Hatakeyama, 2017; George *et al*., 2018). TRIM ligases are characterized by an N-terminal RING ligase domain, zinc-finger B-Box domain(s), and coiled-coil multimerization domain (Meroni and Diez-Roux, 2005; Short and Cox, 2006). A variable C-terminal structure defines nine distinct classes of TRIM proteins and is often considered responsible for providing substrate specificity. Class I TRIMs contain a Cos domain that in some cases binds microtubules (MTs) (Short and Cox, 2006; Cox, 2012), a FN3 domain, and a SPRY domain. In vertebrates, six Class I TRIMs are divided into three paralog pairs, indicative of multiple gene duplication events (Short and Cox, 2006). In contrast, invertebrates have one or two class I TRIM proteins (Boyer *et al*., 2018).

Vertebrate Class I TRIM pairs include TRIM1 and TRIM18, TRIM36 and TRIM46, and TRIM9 and TRIM67. Mutations in TRIM1 (MID2) and TRIM18 (MID1) are associated with X-linked disorders such as intellectual disability and Opitz G/BBB syndrome (Quaderi *et al*., 1997; Cainarca *et al*., 1999; De Falco *et al*., 2003; Geetha *et al*., 2014). TRIM36 regulates dorsal axis formation and cell cycle progression (Cuykendall and Houston, 2009; Miyajima *et al*., 2009). Mutations in TRIM36 are associated with anencephaly (Singh *et al*., 2017). TRIM46 establishes neuronal polarity and the axon initial segment (Van Beuningen *et al*., 2015; Harterink *et al*., 2019). TRIM9 and TRIM67 regulate neuronal morphological changes in response to the axon guidance cue netrin-1 (Winkle *et al*., 2014, 2016; Menon *et al*., 2015; Plooster *et al*., 2017; Boyer *et al*., 2020). TRIM9 localizes to Parkinsonian Lewy bodies and SNPs in TRIM9 may be associated with atypical psychosis (Tanji *et al*., 2010; Kanazawa *et al*., 2013). Paraneoplastic neurological syndromes and small-cell lung carcinoma are associated with TRIM9, TRIM67, and TRIM46 (van Coevorden-Hameete *et al*., 2017; Do *et al*., 2019). These findings demonstrate a role for Class I TRIMs in neuronal form and function.

Loss of the single class I TRIM *dTrim9* in *D. melanogaster* and *madd-2* in *C. elegans* leads to axon branching and guidance defects (Hao *et al*., 2010; Morikawa *et al*., 2011). Similar phenotypes occur with loss of netrin/unc-6 and its receptor, Frazzled/unc-40 (Kolodziej *et al*., 1996; Mitchell *et al*., 1996; Norris and Lundquist, 2011). Similarly, in vertebrates, loss of netrin-1 and its receptor, *deleted in colorectal cancer* (DCC) results in midline crossing defects such as agenesis of the corpus callosum (Serafini *et al*., 1996; Fazeli *et al*., 1997; Bin *et al*., 2015; Yung *et al*., 2015). dTrim9 and MADD2 interact with netrin receptors, Frazzled and UNC40 (Hao *et al*., 2010; Morikawa *et al*., 2011). Analogously, vertebrate TRIM9 and TRIM67 interact with DCC (Winkle *et al*., 2014; Boyer *et al*., 2018). However, unlike loss of *Ntn1* or *DCC*, loss of murine *Trim9* results in corpus callosum thickening, due at least partially to aberrant axon branching within the structure (Winkle *et al*., 2014). In contrast, loss of murine *Trim67* results in thinning of the callosum (Boyer *et al*., 2018). These data suggest that both TRIM9 and TRIM67 function in axon projection downstream of netrin.

We have found that TRIM9 and TRIM67 mediate morphological changes downstream of netrin and DCC by regulating cytoskeletal dynamics and exocytosis (Winkle *et al*., 2014; Menon *et al*., 2015; Plooster *et al*., 2017; Urbina *et al*., 2018; Boyer *et al*., 2020). TRIM9 and TRIM67 interact with and modulate ubiquitination of the actin polymerase VASP to regulate growth cone dynamics and netrin-dependent axonal turning (Menon *et al*., 2015; Boyer *et al*., 2020). TRIM9 mediated ubiquitination of DCC is netrin-1 sensitive and regulates signaling, exocytosis, and axon branching. TRIM9 and TRIM67 regulate the frequency and mode of exocytosis, via interactions with exocytic t-SNAREs SNAP25 and SNAP47, respectively (Winkle *et al*., 2014; Urbina *et al*., 2018, 2020). TRIM9 and TRIM67 continue to modulate neuronal form and function in the mature animal. TRIM9 regulates morphogenesis of adult-born neurons including dendritic arborization, spine density, and localization in the hippocampus (Winkle *et al*., 2016). Loss of *Trim67* results in hypotrophy of the hippocampus (Boyer *et al*., 2018). Along with these anatomical phenotypes, mice lacking *Trim9* and *Trim67* exhibit spatial learning and memory deficits (Winkle *et al*., 2016; Boyer *et al*., 2018), phenotypes typically associated with hippocampal dysfunction.

Together these studies indicate TRIM9 and TRIM67 are important in orchestrating neuronal shape changes, yet the complete repertoire of their function is not known. Most E3 ubiquitin ligases ubiquitinate multiple substrates (Deshaies and Joazeiro, 2009), yet only a few substrates for TRIM9 and TRIM67 have been identified. Further, TRIM9 and TRIM67 interact with proteins without altering their ubiquitination. Therefore, identifying other potential interactors and substrates is critical to understanding the role of TRIM9 and TRIM67 in neuronal development. Here we utilized an unbiased proximity labeling proteomics approach (Roux *et al*., 2012) to identify candidate interacting partners and substrates. This yielded a number of high-priority interaction candidates poised to regulate the MT and actomyosin cytoskeletons, synaptic structure and maintenance, membrane remodeling, and cytosolic transport, processes which are critical to neuronal form and function. We validated the interaction of a subset of candidates with TRIM9 and TRIM67. Analysis of the validated interaction partner Myo16 showed that loss of this unconventional myosin affected axonal branching in a TRIM9 and netrin-1 dependent manner.

## RESULTS

### Proximity dependent labeling identifies interaction partners of TRIM9 and TRIM67

Unbiased identification of ligase interaction partners and substrates is difficult via standard co-immunoprecipitation techniques due to the transient nature of E3 ligase-substrate interactions (Iconomou and Saunders, 2016) and low abundance of ubiquitinated proteins. Proximity-dependent labeling approaches are an effective alternative strategy (Coyaud *et al*., 2015; Cloer *et al*., 2018). To identify candidate TRIM9 and TRIM67 interacting partners and substrates, we employed proximity-dependent labeling with the promiscuous biotin ligase (BirA*) attached to either TRIM9 or TRIM67. This approach exploits biotinylation of proteins within 10 nm of BirA* (Roux *et al*., 2012; Kim *et al*., 2014) and allows identification of transient interaction partners and substrates (**Figure 1A**). Further, we attached a Myc-BirA* to TRIM9 and TRIM67 lacking ligase domains (TRIM9ΔRING and TRIM67ΔRING) to minimize enrichment of biotinylated ubiquitin.

**Figure 1:**
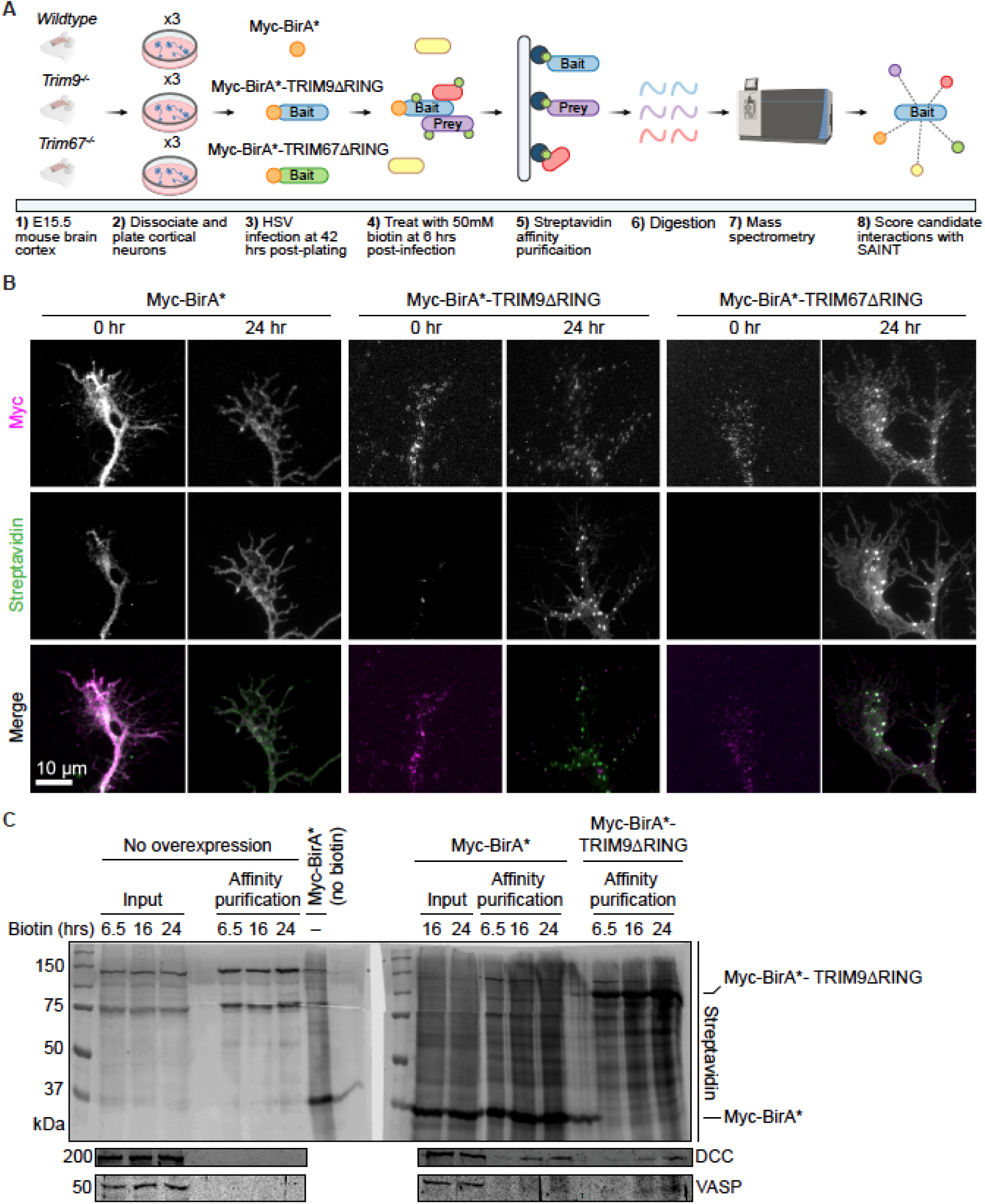
BioID approach to identify candidate interaction partners of TRIM9 and TRIM67 in embryonic cortical neurons. **(A)** Graphical representation of the BioID approach. E15.5 cortical neurons were transduced with HSV carrying Myc-BirA*, Myc-BirA*TRIM9ΔRING, or Myc-BirA*TRIM67ΔRING, media was supplemented with 50 μM biotin. Following cell lysis, biotinylated proteins were affinity purified, enriched proteins were subjected to on-bead trypsinization. Candidate peptides were identified by mass spectrometry. **(B)** Images of axonal growth cones from control Myc-BirA*, Myc-BirA*-TRIM9ΔRING, Myc-BirA*-TRIM67ΔRING expressing neurons at 0 and 24 hrs post-Biotin addition, stained with fluorescent streptavidin and anti-Myc antibody. Merged images show colocalized streptavidin and Myc signals. **(C)** Western blots of streptavidin affinity purification of biotinylated proteins from cortical neurons expressing Myc-BirA* or Myc-BirA*-TRIM9ΔRING. Samples include inputs and affinity purified samples from controls that do not express BirA* (6.5, 16, 24 hrs), input from neurons expressing Myc-BirA* but not supplemented with biotin, inputs (16, 24 hrs) and affinity purified samples (6.5, 16, 24 hrs) from neurons expressing Myc-BirA* and supplemented with biotin, affinity purified samples from neurons expressing Myc-BirA*-TRIM9ΔRING (6.5, 16, 24 hrs). Blots were probed for streptavidin, DCC, VASP.

Since TRIM9 and TRIM67 are enriched in neurons, identification of interaction partners in neurons was critical. MycBirA* constructs were introduced to embryonic cortical neurons by Herpes Simplex Virus (HSV) transduction. To ensure proper localization and function of constructs, we examined localization in neurons and biotinylation of known interaction partners. Myc-BirA*TRIM9ΔRING and Myc-BirA*TRIM67ΔRING localized to the axonal growth cone, primarily to the tips of filopodia and along the periphery of the growth cone (**Figure 1B**), indistinguishable from the localization of endogenous TRIM9 and TRIM67 (Winkle *et al*., 2014; Boyer *et al*., 2018, 2020). In contrast, Myc-BirA* exhibited an expected cytosolic distribution. In neurons supplemented with biotin, the TRIM9ΔRING and TRIM67ΔRING puncta (Myc-stained) overlapped with streptavidin signal (**Figure 1B**), indicating the BirA* was biotinylating proximal proteins. Previously identified TRIM9 substrates, VASP and DCC, were increasingly enriched by affinity purification from lysates of *Trim9*^*-/-*^ neurons expressing Myc-BirA*TRIM9ΔRING after 6, 16, and 24 hrs of biotin addition (**Figure 1C**). In contrast, VASP and DCC enrichment did not increase with biotin treatment duration with Myc-BirA* expression. These results confirmed the feasibility of using an in vitro BioID approach in developing neurons for the identification of candidate E3 ligase substrates and/or interacting partners.

### Potential TRIM9 and TRIM67 interaction candidates affect cytoskeletal change, protein transport, and synaptic structure

Biotinylated proteins were affinity purified from wildtype embryonic cortical neurons expressing Myc-BirA* (negative control), *Trim9*^*-/-*^ neurons expressing Myc-BirA*TRIM9ΔRING, and *Trim67*^*-/-*^ expressing Myc-BirA*TRIM67ΔRING and analyzed using mass spectrometry. 2012 and 2315 total potential interaction candidates were identified for TRIM9 and TRIM67, respectively (**Supplementary Table 1, 2**). The included proteins have a false discovery rate (FDR) ranging from 0.000 (high confidence hits) through 0.7559 (low confidence hits) based on enrichment over MycBirA* hits and variability across replicates. 149 and 151 of all TRIM9 and TRIM67 interaction candidates respectively were classified as high confidence interaction hits with FDR values ≤ 0.25 (**Figure 2A-C**). Of these interaction candidates, 91 were common to both TRIM9 and TRIM67 (**Figure 2C**), approximately 60%. Known interaction partners, including all Enah/VASP family proteins: Enah (TRIM9 interaction FDR: 0, TRIM67 interaction FDR: 0), VASP (TRIM9 interaction FDR: 0, TRIM67 interaction FDR: 0) and Evl (TRIM9 interaction FDR: 0, TRIM67 interaction FDR: 0.0176), and the netrin receptor DCC (TRIM9 interaction FDR: 0.7559, TRIM67 interaction FDR: 0.7559) (Winkle *et al*., 2014; Menon *et al*., 2015; Boyer *et al*., 2018, 2020) were identified by the BioID approach.

**Figure 2:**
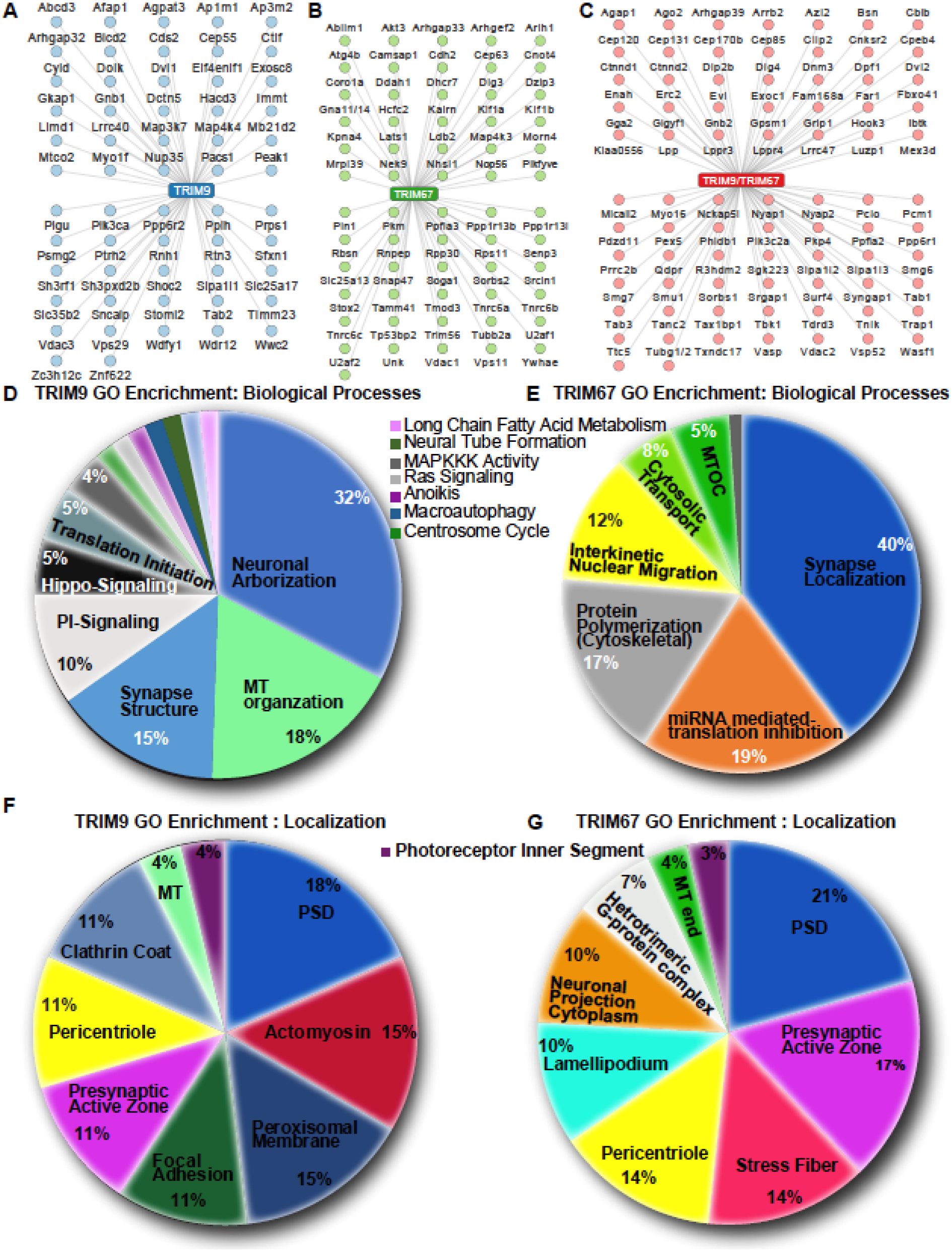
Candidate TRIM9 and TRIM67 interaction partners. **(A-C)** Potential high-confidence (FDR ≤ 0.259) interaction partners unique to TRIM9 (A) unique to TRIM67 (B) or shared (C). **(D,E)** Pie-charts of biological processes enriched via Gene Ontology (GO) classification of high-confidence TRIM9 and TRIM67 interaction partners, respectively. **(F,G)** Pie-charts of cellular localization terms enriched via GO classification of high-confidence TRIM9 and TRIM67 interaction partners, respectively. Percentages for a particular GO term in D-G represent the number of gene IDs that hit against the total number of gene IDs annotated.

Gene Ontology (GO) analysis of high confidence candidates was performed using the ClueGO application in Cytoscape_3.7.2 (Shannon *et al*., 2003; Bindea *et al*., 2009). This approach identified enrichment of protein categories based on function (**Figure 2D,E**) and cellular localization (**Figure 2F,G**). Consistent with findings that TRIM9 regulates morphogenesis, TRIM9 interaction candidates included proteins regulating neuron projection arborization and localizing to the actomyosin and MT cytoskeletons, and localization to clathrin coats, suggesting a possible role in endocytosis (**Figure 2D,F**). Similarly, TRIM67 interaction candidates included proteins that regulated cytoskeletal polymerization and localized to cytoskeletal rich structures, including stress fibers, lamellipodium, the pericentriole, and neuronal projections (**Figure 2E,G**). Although the proteomic analysis was performed in developing neurons prior to synaptogenesis, GO categorization for both TRIM9 and TRIM67 enriched for proteins that mediated localization of proteins to the synapse, were required for maintenance of synapse structures, and localized to pre- and post-synaptic densities. This analysis suggested novel roles for TRIM9 and TRIM67 in signal transduction cascades and protein translation. For TRIM9 this included phosphatidylinositol-mediated signaling, hippo signaling, MAPKKK activity, anoikis, and translation initiation. Similarly, TRIM67 interaction candidates were enriched in cytosolic transport, mRNA mediated inhibition of translation, interkinetic nuclear migration, and MAPKKK activity.

### Validation of TRIM9 and TRIM67 interaction candidates

We focused validation efforts on candidates implicated in actin and MT regulation, membrane remodeling, and motor-based transport. Some high FDR candidates were included in validation efforts since N-terminal tagging of BirA* may have affected the biotinylation of true C-terminal interacting partners, as previously observed (Redwine *et al*., 2017). Indeed the netrin-1 receptor DCC, which interacts directly with the C-terminal SPRY domain of TRIM9 (Winkle *et al*., 2014), and likely the SPRY domain of TRIM67 by homology, exhibited high FDR values (0.7559 for both TRIM9 and TRIM67, 1.05 and 1.26 fold enrichment over control in the TRIM9 and TRIM67, respectively). To validate interactions, we used co-immunoprecipitation (Co-IP) assays of Myc-tagged TRIMΔRING proteins and GFP- or HA-tagged candidate interactors co-expressed in HEK293 cells. This approach overcame issues of transient interactions, low abundance, and unvalidated antibodies to endogenous proteins.

Coro1a (TRIM9 FDR: 0.0027, and TRIM67 FDR: 0) is a type I coronin, a well characterized actin regulator, and modulator of the signaling endosome (Suo *et al*., 2014, 2015; Martorella *et al*., 2017). Coro1a interacted with TRIM67ΔRING, but not TRIM9ΔRING (**Figure 3A**). Sipa1l1, commonly called SPAR [(Spine-Associated Rap GTPase activating protein), (TRIM9 FDR: 0.0082, TRIM67 FDR: 0.5652)], modulates actin dynamics in dendritic spines (Pak *et al*., 2001; Pak and Sheng, 2003; Hoe *et al*., 2009). Sipa1l1 interacted with both TRIM9ΔRING and TRIM67ΔRING, albeit with a higher affinity for TRIM67ΔRING (**Figure 3B**). Myo16 (TRIM9 FDR: 0.0855, TRIM67 FDR: 0.0855), is an unconventional myosin enriched in the developing brain that regulates actin dynamics via the Wave Regulatory Complex (WRC), Arp2/3, and PI3K signaling (Patel *et al*., 2001; Yokoyama *et al*., 2011). FDR values suggested that Myo16 potentially interacted with TRIM9ΔRING and TRIM67ΔRING similarly; co-IP showed that Myo16 interacted more with TRIM67 (**Figure 3C**).

**Figure 3:**
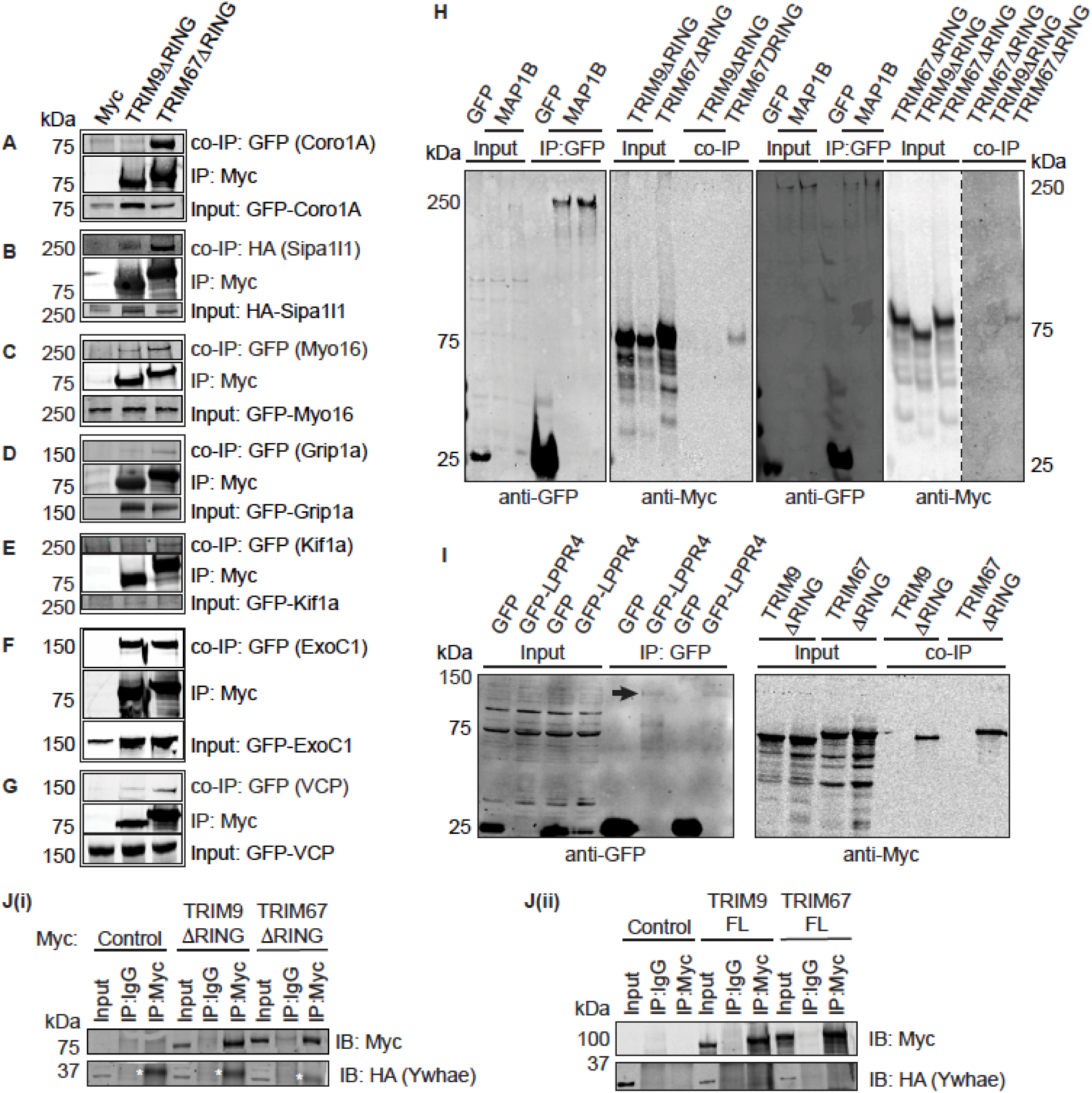
Validation of potential TRIM9 and TRIM67 interaction partners via co-immunoprecipitation assays. **(A-G)** Immunoblots showing co-immunoprecipitation (Co-IP) of GFP- or HA-tagged proteins when immunoprecipitated using an anti-Myc antibody to enrich for Myc-TRIM9ΔRING or Myc-TRIM67ΔRING. (A: Coro1A, B: Sipa1l1, C: Myo16, D: Grip1a, E: Kif1a, F: ExoC1, G: VCP). A-G show blots for inputs for the co-immunoprecipitated proteins, the immunoprecipitated Myc-TRIM9ΔRING or Myc-TRIM67ΔRING and co-immunoprecipitated proteins. **(H)** Immunoblot demonstrating co-IP of Myc-TRIM67ΔRING, but not Myc-TRIM9ΔRING with GFP-MAP1B. Two representative blots are included, one with GFP-control co-expressed with Myc-TRIM9ΔRING and one with GFP-control co-expressed with Myc-TRIM67ΔRING. **(I)** Immunoblot showing Myc-TRIM9ΔRING and Myc-TRIM67ΔRING co-immunoprecipitating with GFP-LPPR4 (PRG1). The black arrow in the blot probed for GFP shows a faint GFP-LPPR4 band. Note that GFP-LPPR4 was not detected in inputs. **(J(i), (ii))** Immunoblots demonstrating HA-Ywhae does not co-IP with Myc-TRIMΔRING or full-length constructs of TRIM9 and TRIM67. The white asterisks in J(i) denote a non-specific band observed in all anti-Myc IP lanes.

Glutamate receptor interacting protein 1 (GRIP1, TRIM9 FDR: 0, TRIM67 FDR: 0) is a kinesin adaptor protein required for homeostatic AMPA receptor trafficking (Setou *et al*., 2002; Hoogenraad *et al*., 2005; Tan *et al*., 2015; Twelvetrees *et al*., 2019). Grip1 interacted with both TRIM9ΔRING and TRIM67ΔRING (**Figure3D**). Kif1a (TRIM9 FDR: 0.271, TRIM67 FDR: 0.0587), a member of the kinesin 3 family, is a MT-based motor protein essential for the transport of synaptic vesicle precursors (Okada *et al*., 1995; Lo *et al*., 2011; Karasmanis *et al*., 2018; Guedes-Dias *et al*., 2019). Kif1a interacted more with TRIM67ΔRING than TRIM9ΔRING (**Figure 3E**). ExoC1 (yeast homolog, Sec3, TRIM9 FDR: 0.0855, TRIM67 FDR: 0), a member of the octameric exocyst complex that plays a critical role in tethering vesicles to the plasma membrane prior to exocytosis (Wu and Guo, 2015; Kampmeyer *et al*., 2017; Yue *et al*., 2017; Liu *et al*., 2018), interacted similarly with TRIM9ΔRING and TRIM67ΔRING (**Figure 3F**). Valosin containing protein (VCP, TRIM9 FDR: 0.7599, TRIM67 FDR: 0.7599) is a clathrin-binding protein and member of the AAA+ ATPase family (Pleasure *et al*., 1993; Ritz *et al*., 2011). VCP interacted with both TRIM9ΔRING and TRIM67ΔRING, but with a higher affinity for TRIM67ΔRING (**Figure 3G**).

MAP1B (TRIM9 interaction FDR: 0.7599, TRIM67 interaction FDR: 0.7599), a large MT-associated protein that plays a role in MT dynamics, neurite branching, and dendritic spine development (Ma *et al*., 2000; Li *et al*., 2006; Tortosa *et al*., 2011; Tymanskyj *et al*., 2012; Bodaleo *et al*., 2016), interacted with TRIM67ΔRING, but not TRIM9ΔRING (**Figure 3H**). Two separate blots are shown to demonstrate the specificity of the interaction with TRIM67; one controlled for TRIM9ΔRING and one for TRIM67ΔRING interaction. LPPR4 (TRIM9 interaction FDR: 0, TRIM67 interaction FDR: 0.0738), commonly called PRG1, is a member of the lipid phosphate phosphatase family. It is a transmembrane protein that functions as an ecto-enzyme and promotes axonal growth by locally depleting lipid phosphates (Bräuer *et al*., 2003; Tokumitsu *et al*., 2010; Petzold *et al*., 2016). GFP-LPPR4 was only detected on western blots after immunoprecipitation (**Figure 3I**). Myc-TRIM9ΔRING and Myc-TRIM67ΔRING were enriched in the GFP-LPPR4 immunoprecipitate, but not the GFP control (**Figure 3I**). Ywhae (TRIM9 FDR: 0.5696, TRIM67 FDR: 0.2572), also called 14-3-3 epsilon, is brain-enriched and required for neurogenesis, neuronal differentiation, and migration (Toyo-Oka *et al*., 2003, 2014). A microduplication of the Ywhae gene locus is associated with autism spectrum disorders (Bruno *et al*., 2010). Ywhae did not interact with full length or ΔRING variants of TRIM9 or TRIM67 (**Figure 3Ji,ii**). These co-IP experiments validated most (7 out of 8) of the candidates tested as interaction partners, supporting the validity of the BioID approach for defining the interactome and suggesting that additional valid interaction partners await confirmation.

Live cell Total Internal Reflection Fluorescence (TIRF) microscopy was performed to examine the colocalization of a subset of validated candidates at the basal surface of developing neurons. This revealed that GFP-Myo16 colocalized with tagRFP-TRIM67ΔRING along the axon and at filopodia tips (**Figure 4A**, arrowheads). Similarly, GFP-LPPR4 colocalized with tagRFP-TRIM67 along filopodia tips along the axon and growth cone (**Figure 4B, arrowheads**). Other interaction partners need analysis to investigate colocalization.

**Figure 4:**
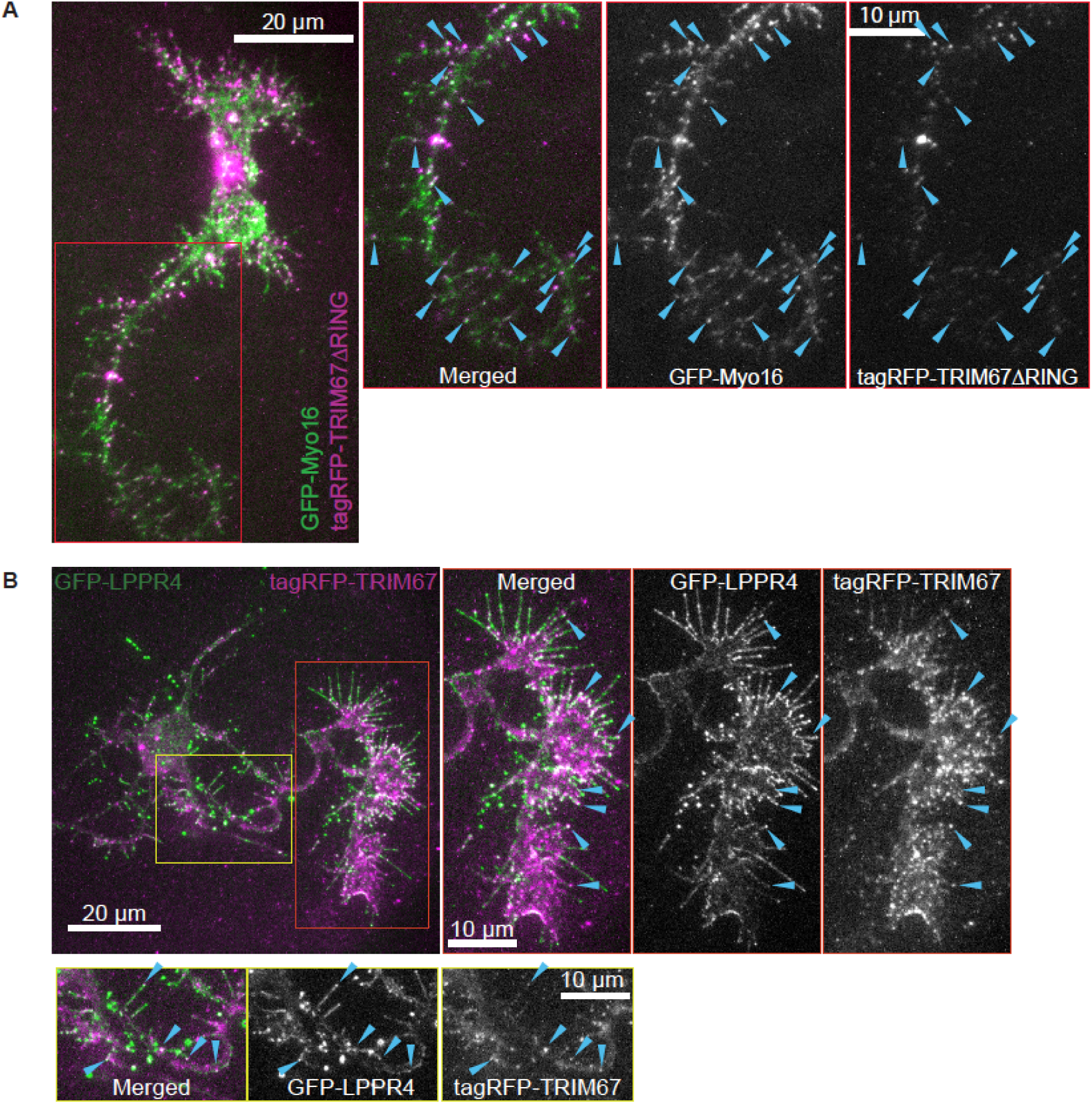
Myo16 and LPPR4 colocalize with TRIM67. **(A)** Representative images to demonstrate colocalization of GFP-Myo16 with tagRFP-TRIM67ΔRING. Zoomed inset and the blue arrowheads show colocalization along the axon and in filopodia along the axon. **(B)** Representative images to demonstrate colocalization of GFP-LPPR4 with tagRFP-TRIM67. Zoomed inset and the blue arrowheads show colocalization along the axon (yellow box) and at the axonal growth cone (red box).

### The Myo16 interaction is unique to TRIM9 and TRIM67

Myo16 expression peaks during neuronal development (Patel *et al*., 2001) where it regulates presynaptic organization in cerebellar Purkinje cells (Roesler *et al*., 2019). TRIM9 and TRIM67 are also expressed in the cerebellum and were recently identified as antibody targets in paraneoplastic cerebellar degeneration (Berti *et al*., 2002; Boyer *et al*., 2018; Do *et al*., 2019). Myo16 localizes along the length of the axon and to the tips of filopodia in hippocampal neurons (Patel *et al*., 2001). Similarly, endogenous TRIM9 and TRIM67 localize to the tips of filopodia, to the periphery of the axonal growth cone, and along the edges of the axon (Winkle *et al*., 2014; Boyer *et al*., 2018, 2020). Similar localization and expression patterns of Myo16, TRIM9, and TRIM67 motivated further investigation of the interaction with Myo16. To confirm that the interaction with Myo16 was specific to TRIM9 and TRIM67, we performed additional co-IP assays. Class I TRIMs share domain architecture; of the three class I paralog pairs, TRIM1 and TRIM18 (MID1) share closest homology with TRIM9 and TRIM67 (Short and Cox, 2006; Carthagena *et al*., 2009). Whereas GFP-Myo16 precipitated with full-length Myc-TRIM9 and Myc-TRIM67, Myo16 failed to precipitate with Myc-TRIM18 (**Figure 5A**). Domain mutants of TRIM67 were employed to map the Myo16 interaction. The FN3 domain of TRIM67 was necessary for mediating TRIM67/Myo16 interaction (**Figure 5B**). The FN3 domain of TRIM9 and TRIM67 are 75% identical, whereas they share less than 30% identity to the FN3 domain of other class I TRIMs (**Supplemental Figure 1A**).

**Figure 5:**
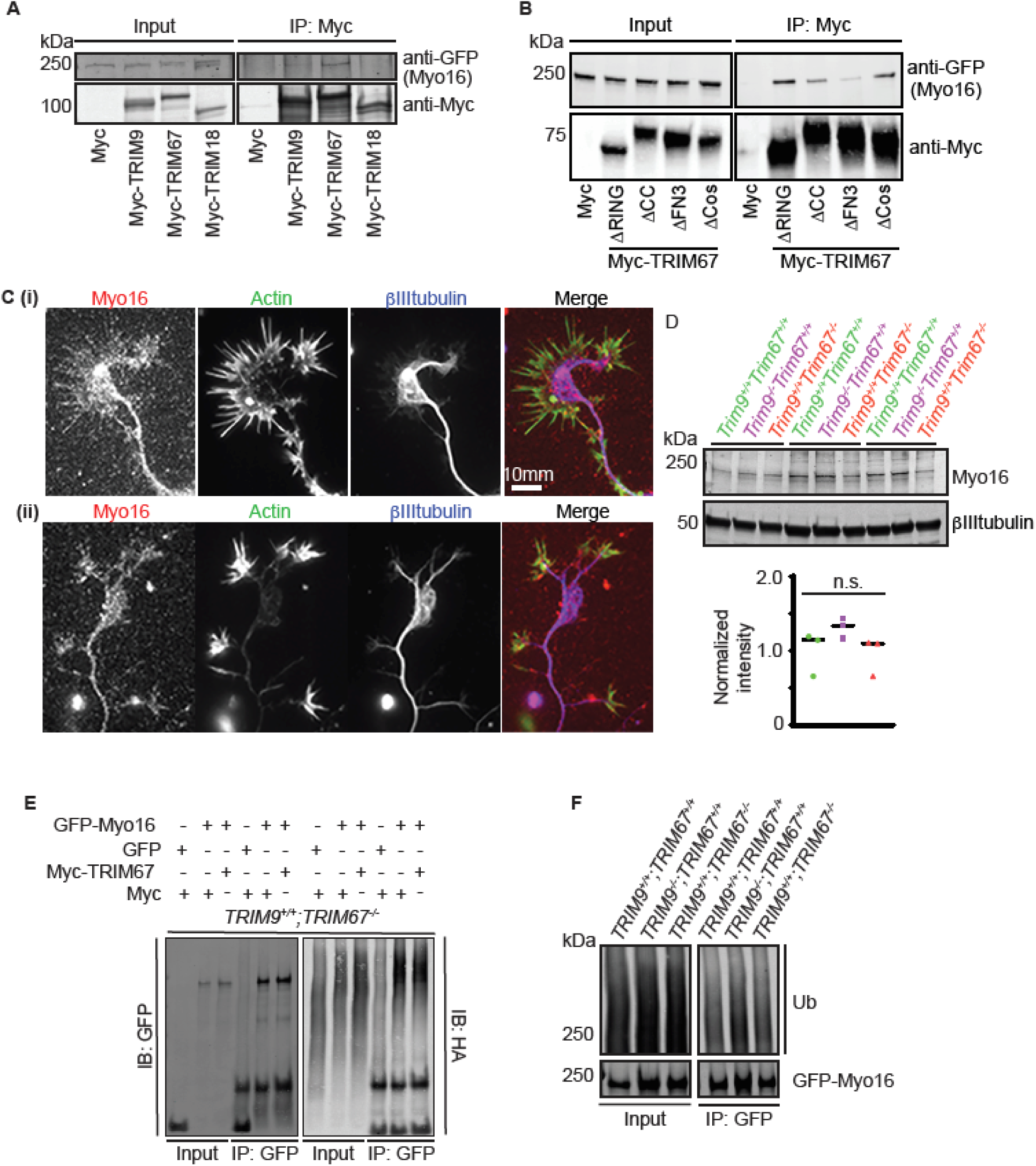
Myo16 interacts specifically with TRIM9 and TRIM67. **(A)** Immunoblot demonstrating GFP-Myo16 co-IP specifically with Class I TRIM proteins, TRIM9 and TRIM67, but not TRIM18 (TRIM proteins used in this assay are full-length and Myc-tagged). **(B)** Representative immunoblot of TRIM67 domain deletion constructs (TRIM67ΔRING, TRIM67ΔCC, TRIM67ΔFN3, TRIM67ΔSPRY) IP demonstrating that the FN3 domain of TRIM67 is necessary for GFP-Myo16 and Myc-TRIM67 interaction. **(C)** Representative images demonstrating endogenous localization of Myo16 in embryonic cortical neurons co-stained with phalloidin for filamentous actin and βIII tubulin to detect MTs. Myo16 localizes to the tips of filopodia and along the axon. **(D)** Immunoblot showing endogenous Myo16 protein levels were not significantly different in cultured embryonic cortical neurons from wildtype (*Trim9*^*+/+*^:*Trim67*^*+/+*^), *Trim9*^*-/-*^ :*Trim67*^*+/+*^, *Trim9*^*+/+*^:*Trim67*^*-/-*^ at 2 DIV. Quantification from 3 biological replicates. **(E)** Representative immunoblot demonstrating GFP (control) and GFP-Myo16 ubiquitination levels in *TRIM67*^*-/-*^ HEK cells expressing either Myc or Myc-TRIM67. The blot was probed with anti-GFP and anti-HA antibodies to visualize Myo16 and ubiquitin respectively. Myo16 ubiquitination status is not altered. **(F)** Representative immunoblot demonstrating GFP-Myo16 ubiquitination levels in wildtype, *TRIM9*^*-/-*^;*TRIM67*^*+/+*^, *TRIM9*^*+/+*^;*TRIM67*^*-/-*^ HEK cells. The blot was probed with anti-GFP and anti-HA antibodies to visualize Myo16 and ubiquitin respectively. Myo16 ubiquitination status is not altered.

With evidence of the specificity of the Myo16 interaction with TRIM9 and TRIM67, we examined whether loss of these proteins affected Myo16. Immunofluorescence indicated that endogenous Myo16 localizes the axon, growth cone, and growth cone filopodia (**Figure 5C**), as we have previously shown for endogenous TRIM9 and TRIM67. Myo16 exhibited a similar localization pattern in the absence of *Trim9* or *Trim67* **(Supplemental Figure 1B)**. Myo16 protein levels in cortical neurons were not altered upon loss of *Trim9* or *Trim67* (**Figure 5D)**. Since the interaction between TRIM67 and Myo16 appeared stronger, we hypothesized that TRIM67 may alter Myo16 ubiquitination. To test this, we performed denaturing immunoprecipitation-based ubiquitination assays of GFP-Myo16 from HEK293 cells in which *TRIM67* had been removed via CRISPR Cas9 genome editing (Boyer et al., 2020). GFP-Myo16 immunoprecipitation contained high molecular weight ubiquitin reactivity in *TRIM67*^*-/-*^ HEK cells expressing Myc alone or Myc-TRIM67 **(Figure 5E)**. In contrast immunoprecipitation of GFP alone revealed no high molecular weight ubiquitin reactivity. These data indicated that Myo16 was ubiquitinated, but it was not TRIM67 dependent. Similar Myo16 ubiquitination was also observed in HEK293 cells in which *TRIM9* had been deleted **(Figure 5F)**. This further revealed that Myo16-ub levels were not different across the three genotypes. Together these data indicated that the Myo16 interaction with TRIM9 and TRIM67 was specific and mediated by the FN3 domain, but the interaction did not detectably alter Myo16 localization, protein levels, or ubiquitination.

### Myo16 is required for netrin-1 dependent axonal branching

We have previously shown that deletion of *Trim9* or *Trim67* independently disrupt axon branching (Winkle *et al*., 2014; Menon *et al*., 2015; Plooster *et al*., 2017; Boyer *et al*., 2020). In *Trim67*^*-/-*^ neurons, basal levels of axon branching are normal, but do not increase in response to netrin-1 addition. In contrast, in *Trim9*^*-/-*^ neurons, basal levels of axon branching are 2-3 fold higher, but also do not respond to netrin-1. We previously found that introduction of full length TRIM67 into *Trim67*^*-/-*^ neurons rescued netrin-dependent axon branching (Boyer et al., 2020). However, TRIM67ΔFN3, which lacks the Myo16 interaction, failed to rescue netrin dependent branching **(Figure 6A)**, potentially implicating Myo16 in netrin-dependent axon branching. To test this hypothesis, Myo16 siRNA were introduced into wildtype cortical neurons at day in vitro 0 (DIV 0), and Myo16 levels were reduced approximately 60% by 2 DIV (**Figure 6B)**. Similar to *Trim67* deletion, Myo16 knockdown did not alter basal axon branching levels, but abrogated netrin-dependent axonal branching **(Figure 6C-D)**. Axon length and progression of neuronal morphogenesis were not perturbed by Myo16 knockdown **(Supplemental Figure 2)**. These data indicated Myo16 was required for netrin-dependent axon branching. Since TRIM67 is known to interact with the netrin receptor DCC (Boyer et al., 2018), yet there is no evidence that Myo16 interacts with DCC, we surmised that overexpression of Myo16 in *Trim67*^*-/-*^ neurons was unlikely to rescue netrin sensitivity and netrin-dependent axon branching. We however wondered if the exuberant axon branching phenotype of *Trim9*^*-/-*^ neurons (Winkle *et al*., 2014) was dependent upon Myo16. Here we confirmed previous data demonstrating that *Trim9*^*-/-*^ cortical neurons showed elevated axon branching density **(Figure 6C,D)**. Reduction of Myo16 in *Trim9*^*-/-*^ cortical neurons reduced axonal branch density **(Figure 6C,D)**, suggesting Myo16 function was required for the aberrant branching phenotype in these neurons. However, axon branching still failed to increase in response to netrin. These data confirm a novel role for Myo16 during axonal branching and netrin responses.

**Figure 6:**
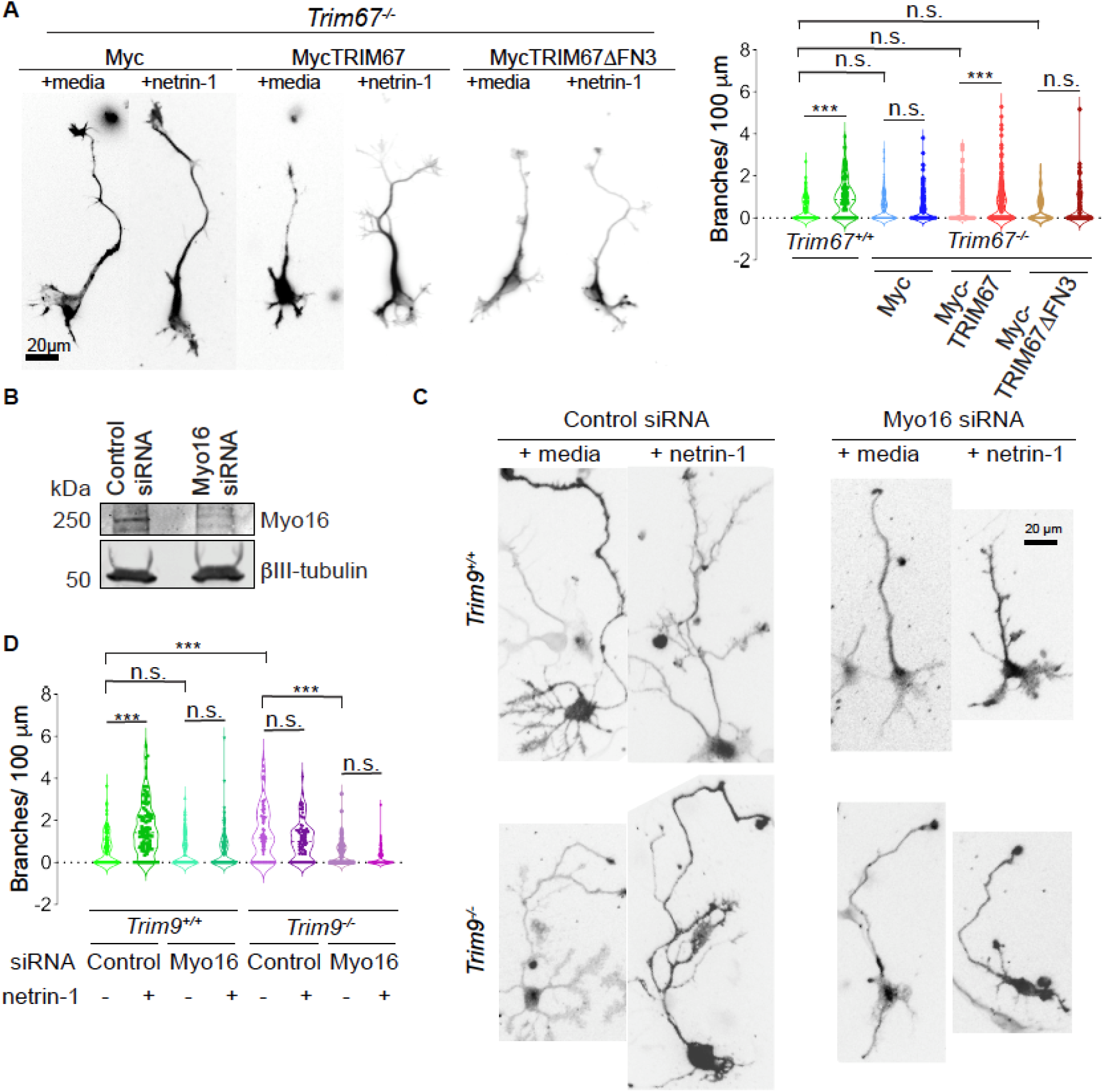
Knockdown of Myo16 inhibits netrin-dependent axonal branching. **(A)** Netrin-1 dependent axon branching assay and quantification in *Trim67*^*-/-*^ neurons expressing Myc, MycTRIM67, or MycTRIM67ΔFN3. The Myc and MycTRIM67 data were published in Boyer et al., 2020. Experiments with MycTRIM67ΔFN3 were performed contemporaneously and not previously published. Staining for filamentous actin (phalloidin) and βIII tubulin were merged, to reveal neuronal morphology. **(B)** Representative immunoblot demonstrating effective knockdown of Myo16 at 2DIV. A prominent Myo16 band that is immunoreactive at approx. 250 kDa in the lane with control siRNA treated lysate shows reduced levels in the lane with Myo16 siRNA treated lysate. **(C)** Representative merged images of GFP (to reveal transfection with control or Myo16 SiRNA) and βIII tubulin (to reveal cell morphology) in wildtype (*Trim9*^*+/+*^) and *Trim9*^*-/-*^ cortical neurons at 3 DIV demonstrate branching phenotypes with or without 24 hrs of netrin treatment. **(D)** Quantification of axonal branching density shown as violin plots with individual data points.

## DISCUSSION

Previously identified interacting partners and substrates suggested that TRIM9 and TRIM67 shared interaction partners, functioned in a yin-yang fashion, and were poised to regulate the cytoskeletal machinery and membrane remodeling machinery that regulate neuronal morphogenesis, function, and connectivity. Here we used proximity ligation to identify previously unidentified candidate interaction partners. This revealed an overlapping interactome of proteins enriched in cellular process regulated by TRIM9 and TRIM67, including neuronal growth and arborization, cytoskeletal dynamics, and membrane remodeling. In addition, novel cellular processes such synapse structure and maintenance were identified as potential TRIM9 and TRIM67 regulation points. We validated a subset of candidates implicated in cytoskeletal dynamics, membrane remodeling, and transport. One validated candidate Myo16, plays a role in netrin-dependent axon branching, potentially in coordination with TRIM9 and TRIM67. Together this study demonstrates that TRIM9 and TRIM67 likely regulate neuronal form and function through a host of interaction partners and potential substrates and this interactome opens up an abundance of new avenues for investigation.

### TRIM9 and TRIM67 regulate the neuronal actin and MT cytoskeletons

Our findings in *Trim9*^*-/-*^ and *Trim67*^*-/-*^ neurons revealed disruptions at multiple stages of neuronal morphogenesis, including filopodia stability, growth cone size and turning, axon branching, dendritic arborization, and dendritic spine density (Winkle *et al*., 2014, 2016; Menon *et al*., 2015; Plooster *et al*., 2017; Boyer *et al*., 2020). GO analysis of candidate interactors here bolster the role of TRIM9 and TRIM67 function in regulating neuronal growth and guidance, with enrichment in neurite outgrowth regulation, and actin and MT cytoskeletal regulation, including actin and MT polymerization, as well as localization to multiple cytoskeletal structures, stress fibers, focal adhesion, MT ends, and lamellipodium. Here we found that the unconventional myosin, Myo16, interacts with both TRIM9 and TRIM67, and plays a critical role in netrin-dependent axon branching. TRIM67 structure/function results and rescue experiments in *Trim9*^*-/-*^ neurons suggested Myo16 regulates netrin-dependent branching in coordination with TRIM9 and TRIM67. The mechanism of this coordinated response is not known. We did find that Myo16 was ubiquitinated, although we did not detect differences in the absence of TRIM9 or TRIM67. However, whether the same residues are ubiquitinated in all genotypes, or whether TRIM9 and TRIM67 redundantly lead to Myo16 ubiquitination is not known. Our previous work found that TRIM9 and TRIM67 interact with each other (Boyer *et al*., 2018) and function in a yin-yang fashion to mediate the non-degradative ubiquitination of VASP (Menon *et al*., 2015; Boyer *et al*., 2020). Non-degradative ubiquitination of VASP was essential for appropriate netrin-1 dependent axonal responses, such as filopodial stability at axonal growth cone and axonal turning. Whether a similar antagonistic relationship exists between TRIM9, TRIM67, and Myo16 is an intriguing hypothesis. Similar questions regarding how other cytoskeletal regulators, like Coro1A and Sipa1l1, are involved in neuronal morphogenesis and regulated by TRIM proteins and downstream of netrin signaling remain to be answered.

Several tubulin isoforms and multiple MT-associated proteins and motors were identified, including MAP1B, DCX, and kinesins Kif1A and Kif1B. Class I TRIMs contain a Cos domain in their C-termini, which has been implicated in MT binding (Short and Cox, 2006). Class I members TRIM18 (MID1) and TRIM46 have both been shown to interact with MTs and regulate MT organization (Van Beuningen *et al*., 2015; Wright *et al*., 2016). As MTs are a critical component in neuronal morphogenesis, investigation into how they are potentially regulated by TRIM9 and TRIM67 is warranted.

### TRIM9 and TRIM67 regulate membrane remodeling

Developing neurons undergo large changes in plasma membrane surface area, driven by exocytosis. Previous work identified that TRIM9 interacts with and sequesters the exocytic t-SNARE SNAP25 from forming SNARE complexes, resulting in constrained constitutive exocytosis and axon branching in developing neurons (Winkle *et al*., 2014). This interaction was netrin sensitive, with netrin treatment increasing SNARE complex formation, exocytosis, and axon branching in a TRIM9-dependent fashion. Strikingly, this increase in exocytosis primarily occurs in the axon, but how the localization of exocytic events occurs is unknown. Here we identified the exocyst component ExoC1 as a validated novel TRIM interactor. Whether this change in the localization of exocytic events involves vesicle tethering, such as the exocyst complex is an intriguing hypothesis.

In contrast to TRIM9, TRIM67 regulates exocytic mode not frequency, promoting full vesicle fusion at the expense of kiss and run fusion (Urbina *et al*., 2020). The t-SNARE SNAP47 was also identified in the proteomic screen described here, specifically as a TRIM67 binding partner. As described elsewhere, we found that SNAP47 was partially responsible for TRIM67-mediated regulation of exocytic mode in developing neurons, but additional proteins must also be involved. Potentially, other membrane-associated proteins like LPPR4 (Sigal *et al*., 2005) or known exocytic regulators like Munc18 and tomosyn, which we identified as candidate TRIM67 binding partners, may also regulate exocytic mode downstream of TRIM67.

### A potential role of TRIM9 and TRIM67 in modulating synapse morphology

Although BioID experiments were performed in neurons prior to synaptogenesis, we found an intriguing enrichment of proteins that localize to the postsynaptic density and the presynaptic active zone and modulate synapse formation and maintenance, such as validated interactors Grip1, Myo16, VASP, PRG1 and Kif1a, as well as yet to be validated candidates such as PSD95, BSN, PCLO, and SynGAP1. We previously demonstrated that deletion of *Trim9* altered dendritic spine density in adult born neurons of the dentate gyrus (Winkle *et al*., 2016), suggesting that TRIM9 may regulate synaptic connectivity. Adult-born hippocampal neurons are critical in spatial learning and memory, which was dramatically deficient in *Trim9*^*-/-*^ mice (Winkle *et al*., 2016). Similarly, genetic deletion of *Trim67* resulted in delayed spatial learning and memory and a loss of cognitive flexibility in the Morris Water Maze task (Boyer *et al*., 2018). Validation of proteins enriched in these categories will give us an insight into how TRIM9 and TRIM67 regulate synapse formation and function. Although how TRIM9 and TRIM67 function at the synapse, and whether this is a developmental role or maintenance role in the adult is unknown, the array of BioID candidates provide multiple pathways to investigate. Many potential scenarios are plausible. This includes regulation of dendritic filopodia formation and spine maturation, based on the role of TRIM9 and TRIM67 in filopodia density and lifetime (Menon *et al*., 2015; Boyer *et al*., 2020). This could be through regulation of VASP or Myo16, both of which have been implicated in spine morphogenesis (Lin *et al*., 2010; Roesler *et al*., 2019). Alternatively, TRIM9 and TRIM67 may play a role in delivery or retrieval of receptors and membrane material (Winkle *et al*., 2014; Urbina *et al*., 2018, 2020) to the pre- or post-synaptic compartments via regulation of proteins such as Grip1, VCP, or Kif1a.

### Other roles for TRIM9 and TRIM67

TRIM67 function is implicated downstream of Ras in neuroblastoma cells (Yaguchi *et al*., 2012). Our studies indicate that both TRIM9 and TRIM67 are involved downstream of netrin and DCC. In the case of DCC, we found that TRIM9 plays a role in signaling. DCC was ubiquitinated in a TRIM9 and netrin-1 sensitive fashion (Plooster *et al*., 2017). This ubiquitination resulted in reduced DCC phosphorylation levels, which constrained binding, autophosphorylation, and activation of focal adhesion kinase (FAK). GO analysis suggested that TRIM9 and TRIM67 may regulate additional signaling networks such as the MAP kinase pathway. Regulation of translation initiation, and miRNA-mediated inhibition of translation are some of the other categories enriched in the BioID candidates. Spatiotemporal regulation of protein translation is critical to the proper form and function of a neuron and regulates axonal branching and pathfinding, memory consolidation and cognitive function (Yoon *et al*., 2009; Buffington *et al*., 2014; Wong *et al*., 2017). Recent evidence also suggests that there are receptor specific interactomes that could serve as a hub for guidance cue induced protein translation, such as in netrin-1/DCC signaling (Koppers *et al*., 2019). Deregulated protein translation has been attributed as the underlying cause of various neurodegenerative disorders (Kapur *et al*., 2017). Validation of these interacting partners remains to be completed. How these pathways are regulated by TRIM9 and TRIM67, and their participation in neuronal morphogenesis and function will be areas of future investigation.

Together previous work along with this study provide evidence that TRIM9 and TRIM67 have multiple overlapping interaction partners and are poised to modulate a number of stages of neuronal morphogenesis, as well as neuronal function. Additional validation and investigations of the TRIM9/TRIM67 interactome are required. However, we propose that these ligases function as molecular rheostats to modulate a shared interactome that coordinates cytoskeletal dynamics and plasma membrane expansion. This coordination facilitates neuronal growth and guidance, essential for establishment of well-connected functional neural networks.

## MATERIALS AND METHODS

### Animals

All mouse lines were on a C57BL/6J background and bred at UNC with approval from the Institutional Animal Care and Use Committee. Timed pregnant females were obtained by placing male and female mice together overnight; the following day was designated as E0.5 if the female had a vaginal plug. Generation of *Trim9*^*-/-*^ and *Trim67*^*-/-*^ mice has been described previously (Boyer et al., 2018; Winkle et al., 2014).

### Cortical neuron culture

E15.5 litters were removed from wildtype, *Trim9*^*-/-*^, and *Trim67*^*-/-*^ pregnant dams by postmortem cesarean section. Dissociated cortical neuron cultures were prepared as described previously (Kwiatkowski *et al*., 2007). Briefly, cortices were micro-dissected, cortical neurons were dissociated with 0.25% trypsin for 20 min at 37°C, followed by quenching of trypsin with Trypsin Quenching Medium (TQM, neurobasal medium supplemented with 10% FBS and 0.5 mM glutamine). After quenching, cortices were gently triturated in neurobasal medium supplemented with TQM. Dissociated neurons were then pelleted at 100 g for 7 min. The pelleted cells were gently resuspended in Serum Free Medium (SFM, neurobasal medium supplemented with B27 (Invitrogen) and plated on coverslips or tissue culture plastic coated with 1 mg/ml poly-D-lysine (Sigma-Aldrich). For axonal length, branching experiments 2 DIV neuronal cultures were treated with bath application of 250 ng/ml netrin-1 for 24 hrs followed by fixation with 4% PFA in PHEM buffer.

### HEK293 cell culture

Wildytpe and *TRIM9*^*-/-*^;*TRIM67*^*+/+*^ HEK293 cells were obtained from Simon Rothenfußer (Klinikum der Universit LJat München, München, Germany) as previously described (Menon et al., 20150). *TRIM9*^*+/+*^;*TRIM67*^*-/-*^ HEK293 cells were generated as previously described (Boyer *et al*., 2020). These cells were grown in DMEM with glutamine (Invitrogen), supplemented with 10% FBS (Hyclone) and maintained at 5%CO_2_/37°C.

### Plasmids, antibodies, and reagents

Plasmids encoding Myc-TRIM9 full-length, Myc-TRIM67 full-length, Myc-TRIM9ΔRING, Myc-TRIM67ΔRING, Myc-TRIM67ΔCC, Myc-TRIM67ΔCos, Myc-TRIM67ΔFN3, Myc-TRIM67ΔCos, Myc-TRIM67ΔSPRY constructs and tagRFP-tagged TRIM67 construct were previously described (Boyer et al., 2018, 2020; Winkle et al., 2014). The pDOWN Gateway entry vectors for expressing Myc-BirA*, Myc-BirA*TRIM9ΔRING or Myc-BirA*TRIM67ΔRING were constructed by VectorBuilder. Herpes Simplex Virus (HSV) vectors for the BirA* constructs were generated by Rachael Neve, MIT proteomics core (currently at Gene Technology Core - Massachusetts General Hospital). The following plasmids were acquired: pDEST307-FLAG-PRG1(LPPR4) (Herbert Geller, NHLBI, NIH; PRG1 was PCR amplied and re-cloned into an eGFP vector, eGFP-PRG1), pML2 (EGFP-N1) - hCoro1A (James Bear, University of North Carolina, Chapel Hill), pEGFP-Myo16 (Anand Srivastava, Greenwood Genetic Center Foundation, VCP(wt)-EGFP (https://www.addgene.org/23971/, originally deposited by Nico Dantuma), Sipa1l1-HA (SPAR-HA) (Daniel Pak, Georgetown University), pEGFP-C3-Sec3 (GFP-ExoC1) (https://www.addgene.org/53755/, originally deposited by Channing Der), GFP-Kif1a (Juan Bonafacino, NICHD, NIH), GRIP1b-EGFP (Rick Huganir, Johns Hopkins University), pcDNA3.1-HA-14-3-3□ (Ywhae) (https://www.addgene.org/48797/, originally deposited by Huda Zhogbi), pCMV-6Myc-hMID1 (Timothy Cox, University of Washington), pEGFP-C1 MAP1B-GFP(Philips Gordon-Week), GFP-hRas CAAX (Richard Cheney, University of North Carolina). Antibodies included goat polyclonal against DCC (Santacruz); rabbit polyclonal against VASP (Santacruz); mouse monoclonal against c-Myc (9E10); mouse monoclonal against mouse β-III-tubulin (Biolegend); mouse monoclonal against HA (12CA5) (Patrick Brennwald, University of North Carolina); rabbit polyclonal against Myo16 (Richard Cameron, Augusta University); rabbit polyclonal against Myo16 (Proteintech, 25104-1-AP); fluorescent secondary antibodies. Other fluorescently labeled reagents included streptavidin and phalloidin labeled with Alexa Fluor 561. Recombinant chicken myc-netrin-1 was concentrated from HEK293 cells (Lebrand et al., 2004; Serafini et al., 1996).

### Proximity dependent biotin identification (BioID), affinity purification and peptide preparation

The negative control Myc-BirA* and Myc-BirA*TRIM9ΔRING (amino acids 132-710) or Myc-BirA*TRIM67ΔRING (amino acids 164-783) constructs were cloned into Herpes Simplex Virus (HSV) vectors using Gateway Cloning (VectorBuilder and Rachael Neve, MIT Viral Vector Core). These vectors were packaged into Short Term Herpes Simplex Virus (HSV) and driven under a IE 4/5 promoter. GFP expression is also driven in tandem downstream of a mCMV promoter. Both promoters are equivalent in the timing of expression and the strength. GFP signal serves as a good indicator of transduction efficiency and timing of expression.

E15.5 cortical neurons from wildtype and *Trim9*^*-/-*^ or *Trim67*^*-/-*^ E15.5 litters were dissociated and plated on PDL (Sigma)coated tissue culture dishes. Approximately 40 hours post-plating neurons were infected with HSV (MOI = 1.0) carrying either the negative control Myc-BirA*, Myc-BirA*TRIM9ΔRING, or Myc-BirA*TRIM67ΔRING. 6 hrs post-infection, cultures were supplemented with 50 μM Biotin for 24 hrs. After incubation, cells were lysed using RIPA buffer (150 mM NaCl, 25 mM Tris-HCl, pH 7.5, 0.1% SDS, 1.0% NP-40, 0.25% Deoxycholic acid, 2 mM EDTA, 10% glycerol, protease and phosphatase inhibitors). Lysates were treated with 0.5 ml Benzonase/ml lysate with end-to-end rotation for 1 hr at 4°C. Lysates were cleared by centrifugation, 16,000xg for 30 min at 4°C. Biotinylated proteins were enriched using Streptavidin-conjugated Sepharose beads (GE Healthcare Life Sciences). Beads were washed once with RIPA buffer, two consecutive washes with TAP lysis buffer (50 mM HEPES (pH 8.0), 10% glycerol, 150 mM NaCl, 2 mM EDTA, and 0.1% NP-40) and three consecutive washes with 50 mM ammonium bicarbonate. Enriched proteins were digested with 2.5 μg trypsin at 37°C for 16 hrs. Peptides were eluted using the *Rapi*GEST SF Surfactant protocol (Waters). Eluted peptides were desalted using C18 columns followed by an ethyl acetate cleanup. Peptides were then stored at -20°C until analysis by mass spectrometry.

### Mass spectrometry and data analysis

Reverse-phase nano-high-performance liquid chromatography (nano-HPLC) coupled with a nanoACQUITY ultraperformance liquid chromatography (UPLC) system (Waters Corporation; Milford, MA) was used to separate trypsinized peptides. Trapping and separation of peptides were performed in a 2 cm column (Pepmap 100; 3-m particle size and 100-Å pore size), and a 25-cm EASYspray analytical column (75-m inside diameter [i.d.], 2.0-m C18 particle size, and 100-Å pore size) at 300 nL/min and 35°C, respectively. Analysis of a 150-min. gradient of 2% to 25% buffer B (0.1% formic acid in acetonitrile) was performed on an Orbitrap Fusion Lumos mass spectrometer (Thermo Scientific). The ion source was operated at 2.4kV and the ion transfer tube was set to 300°C. Full MS scans (350-2000 m/z) were analyzed in the Orbitrap at a resolution of 120,000 and 1e6 AGC target. The MS2 spectra were collected using a 1.6 m/z isolation width and were analyzed either by the Orbitrap or the linear ion trap depending on peak charge and intensity using a 3 s TopSpeed CHOPIN method. Orbitrap MS2 scans were acquired at 7500 resolution, with a 5e4 AGC, and 22 ms maximum injection time after HCD fragmentation with a normalized energy of 30%. Rapid linear ion trap MS2 scans were acquired using an 4e3 AGC, 250 ms maximum injection time after CID 30 fragmentation. Precursor ions were chosen based on intensity thresholds (>1e3) from the full scan as well as on charge states (2-7) with a 30-s dynamic exclusion window. Polysiloxane 371.10124 was used as the lock mass.

For the data analysis all raw mass spectrometry data were searched using MaxQuant version 1.5.7.4. Search parameters were as follows: UniProtKB/Swiss-Prot mouse canonical sequence database (downloaded 1 Feb 2017), static carbamidomethyl cysteine modification, specific trypsin digestion with up to two missed cleavages, variable protein N-terminal acetylation and methionine oxidation, match between runs, and label-free quantification (LFQ) with a minimum ratio count of 2. The mass spectrometry proteomics data have been deposited to the ProteomeXchange Consortium via the PRIDE (Perez-Riverol *et al*., 2019) partner repository with the dataset identifier PXD021758. To rank candidate protein-protein interactions by likelihood of interaction, LFQ values for proteins identified in control and experimental conditions were input into SAINTq (version 0.0.4). Interactions were sorted by SAINT’s interaction probability and a false discovery rate (FDR) threshold was assigned to each of the identified proteins.

### Data representation and Gene Ontology (GO) analysis

Cytoscape _3.7 open source bioinformatics interaction visualization software was used to visualize potential interaction candidates using a Prefuse Force Directed Layout in which the source node (TRIM9, TRIM67, or both) appears in the center of the network. Gene ontology enrichment analysis was performed using the ClueGo application available for download in the Cytoscape_3.7.

### Transfection of cortical neurons and HEK293 cells

For transfection of plasmids and siRNA, dissociated cortical neurons were resuspended in Lonza Nucleofector solution (VPG-1001) and electroporated with an Amaxa Nucleofector according to manufacturer protocol. HEK293 cells were transfected using Polyplus jetPRIME® reagent or Lipofectamine 2000 as per manufacturer’s protocol.

### Co-immunoprecipitation assays

For co-immunoprecipitation assays HEK293 cells were transfected with plasmids expressing tagged versions of the proteins-of-interest and Myc or GFP-tagged TRIM9 or TRIM67 full length or ΔRING constructs or Myc-tagged TRIM67 domain deletion constructs. 16-18 hrs post-transfection the cells were lysed using the immunoprecipitation (IP) buffer (10% glycerol, 1% NP-40, 50 mM Tris pH 7.5, 200 mM NaCl, 2 mM MgCl_2_ and protease and phosphatase inhibitors). Cells were scraped from the dish and transferred into tubes, then centrifuged at 18.4K g for 10 min. Approximately 500–1,000 µg of protein per sample was used per co-IP. Myc-tagged proteins were enriched using anti-Myc (9E10) antibody. Antibody was incubated overnight with the lysate following which Protein A coupled agarose beads were added to the lysate-antibody mix. After 2 hrs of incubation the beads were precipitated, washed twice with IP buffer and then boiled with 2X sample buffer. GFP-Trap beads (Chromotek) were used to enrich GFP-tagged proteins. The beads were incubated with the lysate for 1 hr. The beads were then precipitated, washed twice with IP buffer and then boiled with 2X sample buffer.

### Ubiquitination assay

Wildytpe, *TRIM9*^*-/-*^;*TRIM67*^*+/+*^, *TRIM9*^*+/+*^;*TRIM67*^*-/-*^ HEK cells were transfected with HA-Ub, and GFP-Myo16, and in some cases Myc or Myc-TRIM67, using Lipofectamine 2000 and cultured for 24 hrs. These cells were treated with 10 μM MG132 for 4 hours. The cells were then lysed in IP buffer (20 mM Tris-Cl, 250 mM NaCl, 3 mM EDTA, 3 mM EGTA, 0.5% NP-40, 1% SDS, 2 mM DTT, 5 mM NEM (N-ethylmaleimide), 3 mM iodoacetamide, protease and phosphatase inhibitors pH=7.3-7.4). For 5-6 million cells 270 µl of ubiquitin IP buffer was added and incubated on ice for 10min. Cells were removed from the dish and transferred into tubes. 30 μl of 1X PBS was added and gently vortexed. Samples were boiled immediately for 20 minutes, until clear, then centrifuged at 18.4K g for 10 minutes. The boiled samples were diluted using IP Buffer without SDS to reduce the SDS concentration to 0.1%. Immunoprecipitations were performed under conditions in which only covalent interactions were preserved with anti-GFP (mouse) antibody overnight at 4°C following which Protein A agarose beads (Santa Cruz) were used to precipitate the antibody bound complex.

### Immunoblotting

The immunoprecipitated proteins from co-immunoprecipitation assays or ubiquitination assays or endogenous neuronal lysates were separated on reducing polyacrylamide gels and transferred to nitrocellulose membrane using standard procedures. The blots were probed with tag-specific or protein-specific primary antibodies followed by species-specific far-red conjugated secondary antibodies (Licor). Signal was detected using an Odyssey Imager (Licor).

### Immunofluorescence

Neurons fixed with 4% PFA in PHEM buffer were permeabilized for 10 min in 0.1% Triton X-100. Permeabilized cells were blocked for 30 min in 10% donkey serum, and stained with indicated primary antibodies for 1 hr at RT. The primary antibodies were then washed off (4⨯5 min washed in 1X PBS) following which species-appropriate fluorescently labeled secondary antibodies or fluorescently labeled phalloidin were added and allowed to incubate for 1 hr at RT. After the secondary antibodies/phalloidin was washed off cells were mounted in a TRIS/glycerol/n-propyl-gallate-based mounting medium and stored at 4°C till they were ready to be imaged.

### Colocalization experiments

GFP-tagged Myo16 and tagRFP-TRIM67 constructs were expressed in *Trim67*^*-/-*^ dissociated cortical neurons. Colocalization was observed using Total Internal Reflection Fluorescence (TIRF) microscopy. For live-cell TIRF and time-lapse imaging cells were maintained at 37°C, 5% CO_2_ and humidity using a stage top incubator (Tokai Hit).

### Microscopy and Image acquisition

All immunofluorescence and live-cell images were collected on an Olympus IX81-ZDC2 inverted microscope. The following objective lenses were utilized: a UPLFLN 40×/1.39-NA differential interference contrast (DIC) objective (Olympus), UAPON 100×/1.49-NA DIC TIRF objective (Olympus). The microscope is equipped with an automated XYZ stage (Prior), and an Andor iXonelectron multiplying charge-coupled device. Cells for live imaging were maintained at 37°C and 5% CO_2_ using a TokaiHit on-stage cell incubator. Images were acquired using Metamorph acquisition software. Image analysis was performed in Fiji. Axon branches were defined as extensions off the primary axon (more than two times longer than other neurites) that were ≥20 µm in length.

### Statistics

All experiments were performed in three or more replicates. Nonparametric analysis of variance (ANOVA) with Tukey’s post-hoc correction, Benjamini-Hochberg procedure, or multiple logistic regression was performed for data with multiple comparisons. Mass spectrometry data analysis statistics has been described above. P-values are listed in figures or figure legends (not significant (n.s.) at P > 0.05; *, P < 0.05; **, P < 0.01; and ***, P < 0.005).

## Acknowledgements

We thank Vong Thoon, Caroline Monkiewicz, Carey Hanlin, Janee Cadlett-Jete, and Natalia Riddick for mouse colony management. We thank Emily Cousins for help with troubleshooting the BioID approach and sample submission for mass spectrometric analysis. This work was supported by National Institutes of Health R21MH108970 and R35GM135160 (S.L.G).

## Abbreviations used

BioID: proximity-dependent biotin identification
BirA*: Promiscuous bifunctional ligase/repressor BirA
Co-IP: co-immunprecipitation
Coro1a: coronin, actin binding protein 1A
DCC: deleted in colorectal cancer
ExoC1: exocyst complex component 1
FDR: False Discovery Rate
GO: Gene Ontology
GRIP1: Glutamate receptor interacting protein 1
HSV: Herpes Simplex Virus
KIF1A: Kinesin family member 1A
MAP1B: Microtubule associated protein 1B
MS: Mass Spectrometry
MT: microtubule
Myo16: myosin XVI
PRG-1: phospholipid phosphatase related 4 (Plppr4)
SNAP25: synaptosome associated protein 25
SNAP47: synaptosome associated protein 47
TIRF: Total Internal Reflection Fluorescence
TRIM67: tripartite motif containing 67
TRIM9: tripartite motif containing 9
Ywhae: 14-3-3 epsilon

## SUPPLEMENTAL MATERIAL

**Supplemental Table 1: Candidate TRIM9 and TRIM67 interaction partners**. All candidate TRIM9 (sheet 1) and TRIM67 (sheet 2) interaction partners listed in order of their FDR (lowest to highest).

**Supplemental Table 2: Peptide counts and label free quantitation intensities for all identified proteins**.

## Supplemental Figures and Legends

**Supplementary Figure 1:**
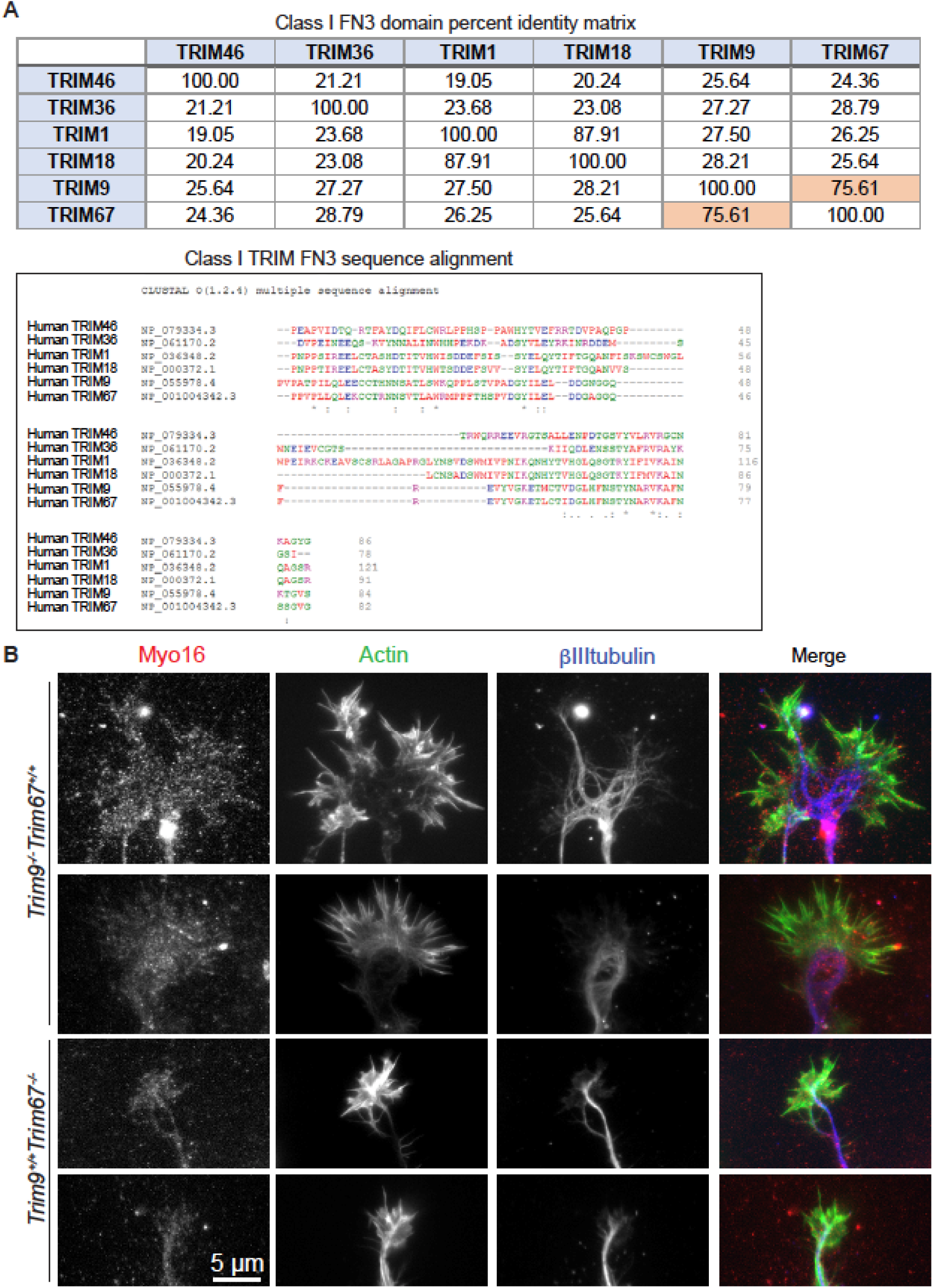
Myo16 interacts specifically with TRIM9 and TRIM67. **(A)** Representative images demonstrating endogenous localization of Myo16 in *Trim9*^*-/-*^ and *Trim67*^*-/-*^ embryonic cortical neurons co-stained with phalloidin for filamentous actin and βIII tubulin to detect MTs. Myo16 localizes to the tips of filopodia and along the axon. **(B)** Sequence analysis and alignment of FN3 domains of human Class I TRIM proteins.

**Supplementary Figure 2:**
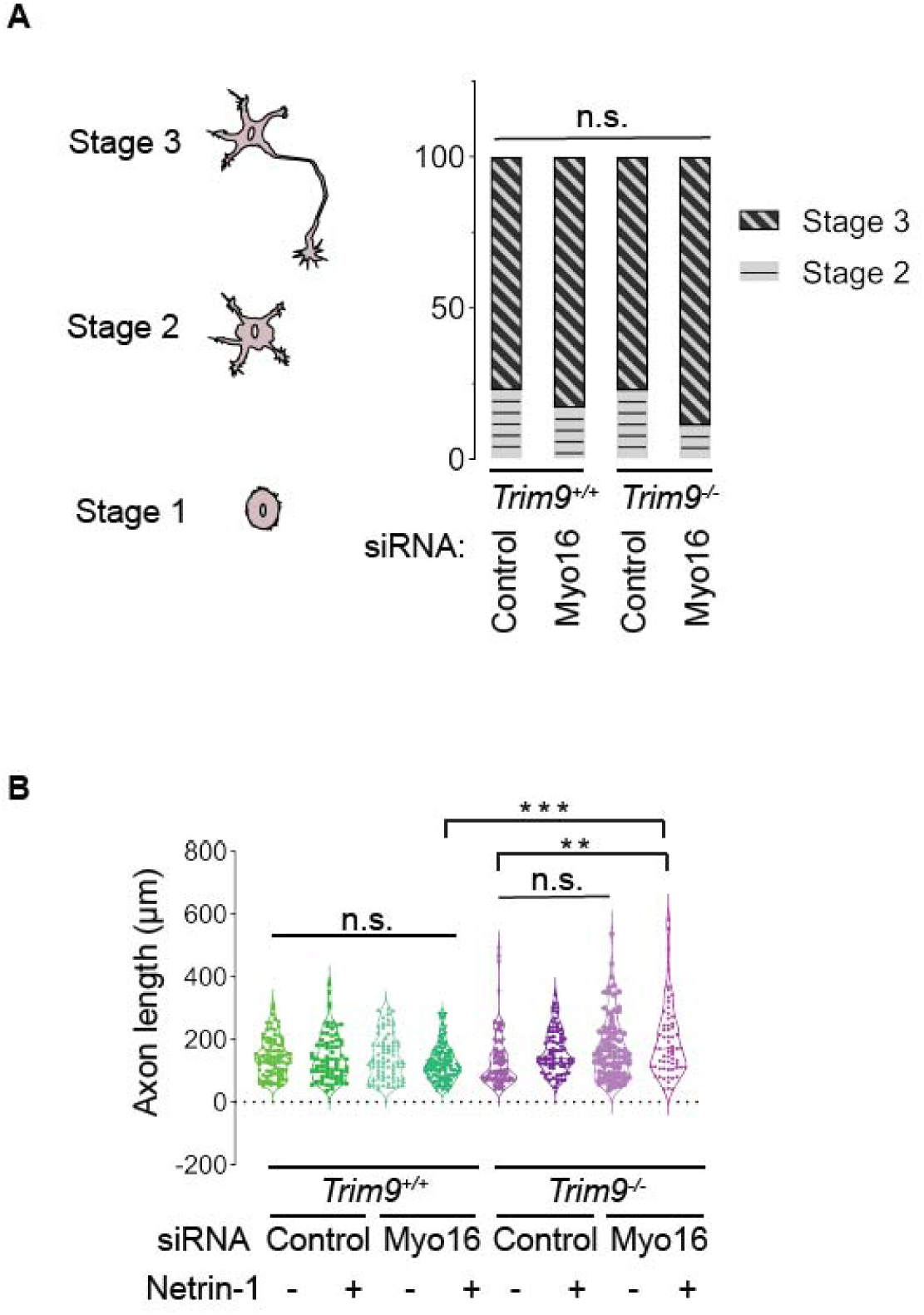
Knockdown of Myo16 inhibits netrin-dependent axonal branching. **(A)** Myo16 knockdown does not alter neuronal staging (stage 2 vs. stage 3) in in-vitro cultures of embryonic cortical neurons. **(B)** Quantification of axon length from cortical neurons at 3 DIV in wildtype (*Trim9*^*+/+*^:*Trim67*^*+/+*^), *Trim9*^*-/-*^:*Trim67*^*+/+*^ with or without 24 hrs of netrin treatment shown as violin plots with individual data points.

## Author contributions

SM: Project design; data acquisition, curation, and validation, data analysis, data interpretation; training undergraduate researchers, manuscript and figure preparation; DG: Mass spectrometry (MS) and MS data analysis; visualization; TH: Myo16 KD analysis; EWC: MS sample quality control and peptide quantification; NPB: *Trim67*^*-/-*^ rescue branching experiment; CH, AJB, ECJ, JA: Co-immunoprecipitation experiments; MBM: Oversaw MS methods and MS data analysis; manuscript editing; SLG: Project conceptualization, design, administration, funding acquisition; interpretation of data ; manuscript and figure preparation and approval.

## Notes

### Competing Interest Statement

The authors have declared no competing interest.

### Summary of Updates

Errors in author names

